# Eastern Joshua tree arbuscular mycorrhizal fungal mycobiomes largely consistent across roots, soils, and seasons

**DOI:** 10.1101/2025.03.05.641720

**Authors:** Arik Joukhajian, Sydney I. Glassman

## Abstract

The Mojave Desert is home to iconic Joshua trees threatened by climate change. Most desert plants form mutually beneficial partnerships with arbuscular mycorrhizal fungi (AMF), yet the AMF of the Eastern Joshua tree (Yucca jaegeriana) remain completely uncharacterized. We tested how Y. jaegeriana AMF spore abundance, richness, and composition varied when sampling 20 trees across 4 seasons from roots versus soils. We confirmed root colonization via staining, assessed spore abundance via microscopy, and used Illumina MiSeq to sequence AMF virtual taxa (VT) with WANDA AML2 primers. We identified 12 spore morphotypes and 47 VTs across 5 families within Glomeromycotina and the most abundant VT Glomus VTX00294 appeared in 87% of soil and root samples. The majority of VTs (26/47) were present across all seasons and were shared among soil and roots (38/47) with more VTs unique to soil. In soil, per tree mean spore abundance and AMF richness was lowest in Summer but consistent across other seasons with richness ranging from 8.8 to 11.5 VTs and mean root richness consistent across seasons. We conclude that sampling from soils rather than roots and any season other than Summer will yield the most diverse AMF communities.

## Introduction

Mycorrhizal fungi enable many desert plants to survive at the edge of climatic extremes, particularly by increasing access to water (Augé 2001; Miriti et al. 2007). Yet, many desert plant species face contraction of their reproductive ranges as climate change exacerbates desert conditions (Guida et al. 2014). The iconic Joshua trees are among the many desert plants facing such threats, and the Western Joshua tree, Yucca brevifolia, has shown both long-term predicted declines and observed range shrinkage in reproductive viability due to climate change (Sweet et al. 2019). Yuccas, like most desert plants, associate with symbiotic arbuscular mycorrhizal fungi (AMF) within the Glomeromycotina (Titus et al. 2002; James et al. 2020) for increased access to nutrients and water (Smith and Read 2010). As parts of the Joshua tree habitat become unsuitable for reproduction, it is critical to understand their microbial associations in the cooler climate refugia where Joshua trees are expected to thrive.

Despite the iconic nature of Joshua trees, many open questions remain about the basic biology of these plants and their associated microbiomes. Yucca jaegeriana, the Eastern Joshua tree, which is densest in cooler climate refugia (Esque et al. 2023), was recently recognized as a distinct species from the Western Joshua tree based on morphology (Lenz 2007), its unique species of obligate pollinator moth (Smith et al. 2008), and genomic sequencing (Royer et al. 2016). AMF have been found in desert ecosystems across the globe (Vasar et al. 2021), including 3 species of Yucca (Titus et al. 2002; Tawaraya 2003; Harrower and Gilbert 2021), yet the AMF of Y. jaegeriana remain completely uncharacterized. One existing study found 37 AMF virtual taxa (VTs, or roughly the equivalent of AMF species based on 18S rRNA sequences (Öpik et al. 2010)), associated with the roots of sister species Y. brevifolia within Joshua Tree National Park (JTNP). AMF taxa in Y. brevifolia roots were clustered phylogenetically by elevation, and experimental seedling inoculation showed that all sources of native AMF inoculum increased plant biomass after six months (Harrower and Gilbert 2021). This suggests that Y. jaegeriana will be similarly colonized with diverse AMF taxa important to its growth and survival.

Beyond being completely uncharacterized within Y. jaegeriana, little is known in general about how AMF vary in richness and composition across seasons or in roots versus soils. The impact of seasons on richness and community composition of AMF can vary from weak to strong in temperate grasslands (Dumbrell et al. 2011; Sepp et al. 2019). Seasonality also had no effect on AM community composition in a temperate forest (Davison et al. 2012) but a strong effect in an arid shrubland and a subtropical forest (Sánchez-Castro et al. 2012; Maitra et al. 2019). Finally, AMF richness remained stable throughout the year in semi-arid shrublands (Chaudhary et al. 2014). Moreover, sample type (root, soil, spore, or rhizosphere) drove significant differences of community composition in grasslands, wheat prairies, agricultural sorghum fields, and a subtropical forest (Hempel et al. 2007; Ellouze et al. 2018; Sepp et al. 2019; Gao et al. 2019), suggesting that sampling root or soil can also lead to differing AMF communities. Therefore, a comprehensive characterization of Y. jaegeriana AMF communities should sample across seasons and sample types.

AMF can also vary in spore abundance and root colonization across seasons. If host plants have different active seasons, then AMF species can show seasonal preferences for sporulation (Bever et al. 2001). Sporulation can also vary based on climate. For example, sporulation trended with precipitation in a water limited semi-arid region in Brazil (Da Costa et al. 2021). However, this seasonality of spore abundance disappeared in a Brazilian rainforest with high humidity (Kemmelmeier et al. 2022). AMF root colonization of the perennial Mojave desert shrubs Larrea tridentata and Ambrosia dumosa showed strong seasonal changes after cool-season rains (Apple et al. 2005) but these seasonal changes in colonization went away during drought (Titus et al. 2002; Clark et al. 2009). Therefore, it is likely that AMF sporulation and colonization would vary across seasons in a desert plant like Y. jaegeriana.

Since AMF species can vary seasonally, there may only be a few AMF species present in the core community year-round. Core microbiomes can either refer to a few taxa repeatedly identified across time within the same host individual, or to core taxa that are shared across all host individuals within a population (Risely 2020). Some evidence points to AMF occurring as core community members. For example, 4 AMF sequence phylotypes formed a temporal core community for two plants in an arid shrubland (Sánchez-Castro et al. 2012), potatoes appear to have a core community of AMF partners across the Andes mountains despite a wide geographic range (Senés-Guerrero and Schüßler 2016), and some AMF species were consistently found with the Western Joshua tree across long physical distances (Harrower and Gilbert 2021).

Dominance within a particular season does not necessarily indicate membership of the core microbiome. For example, 2 AMF taxa that were highly dominant in June disappeared by September in sorghum fields (Gao et al. 2019). Thus, sampling across both time and space is necessary to identify a core microbiome.

Finally, another factor potentially impacting the AMF of Y. jaegeriana is fire. A mix of intermittent drought, exotic grasses (McAuliffe 2016), and Winter rains (Tagestad et al. 2016) primed the Cima Dome, which was home to the densest population of Y. jaegeriana, for the 2020 Dome Fire, killing > 1 million trees (NPS. 2020). High-elevation regions like the Cima Dome are among the 10% of habitat expected to continually support Joshua tree reproduction in the face of ongoing climate change (Sweet et al. 2019) making it a particularly important location to characterize Y. jaegeriana AMF. While AMF are typically more resistant to high temperatures than other soil microbes (Allen et al. 2011), post-fire communities of AMF may be shaped by the succession of their plant hosts (Neuenkamp et al. 2018) and the varying survival rates of initial community members based on their differing spore traits (Hopkins and Bennett 2023).

There may not be any correlation between traits that allow AMF to support early successional plants and traits for fire survivorship, so turnover of AMF following fire is possible.

Here, we comprehensively characterize the Y. jaegeriana AMF community via both traditional microscopy and modern next-generation sequencing methods in root and soils and across seasons. We collected root and soil samples from 20 healthy Eastern Joshua trees from each of the four seasons spanning from June 2021 to March 2022 with 16 trees outside and 4 surviving but scarred trees within the 2020 Dome Fire burn scar. We asked if AMF spore abundance, richness, and composition varied across seasons and sample types (roots versus soil) and if a core community exists across roots/soils and seasons. We further tested if being within a burn patch affected AMF richness and composition. We predicted that spore abundance would significantly increase during seasons of high precipitation, corresponding to Summer and Winter samplings, which would also vary in AMF community composition.

However, we predicted that AMF richness would be lower in Eastern Joshua trees within the burn scar due to reduction of surrounding desert shrubs and grasses. We also predicted that roots would provide greater AMF species richness than soil since Glomeraceae, which were abundant in Western Joshua tree Y. brevifolia (Harrower and Gilbert 2021), preferred roots over rhizosphere or extra-radical mycelium in semi-arid plants (Varela-Cervero et al. 2015; Alguacil et al. 2016). Finally, we predicted that a Y. jaegeriana AMF core community would consist of members of the Glomeraceae and Ambispora from Ambisporaceae, corresponding to taxa that were abundant at a similar elevation band for Y. brevifolia (Harrower and Gilbert 2021).

## Methods

### Sampling

We selected 20 healthy Yucca jaegeriana trees (Fig. 1A) with a minimum of 5 florets and 2m height as our focal plants in Mojave National Preserve, CA (Table S1). Sixteen trees (#1- 16) were selected east of the Morningstar Mine Rd which acted as a barrier for the Dome Fire, while four fire-scarred but surviving trees (#17-20) were selected within the burn perimeter (Fig. 1B). We sampled at the beginning of each season with Summer on June 23, 2021, Fall on September 9, 2021, Winter on December 21, 2021, and Spring on March 30, 2022. We collected and extracted DNA from both roots and surrounding soil for a total of 160 samples (2 types (roots vs soil) X 4 seasons (Summer, Fall, Winter, Spring) x 20 trees). We used releasable bulb planters and wiped off debris with 70% ethanol between plots to collect 2-3 soil cores of the top 10 cm of soil from close to the base of the trunk of each tree at each time point, avoiding previously sampled edges of the base when possible (Fig. 1A). Soil was stored on ice and returned to UC Riverside where it was sieved (2mm) within 24-48 hours of collection, during which we separated roots from within the soil sample. We stored approximately 10g of soil and 1g of roots at -80°C for downstream DNA extraction and air-dried the remaining soil. We stored additional roots in -80°C or in 50% ethanol solution in 4°C for staining and visualization of fungal structures.

**Figure 1.**
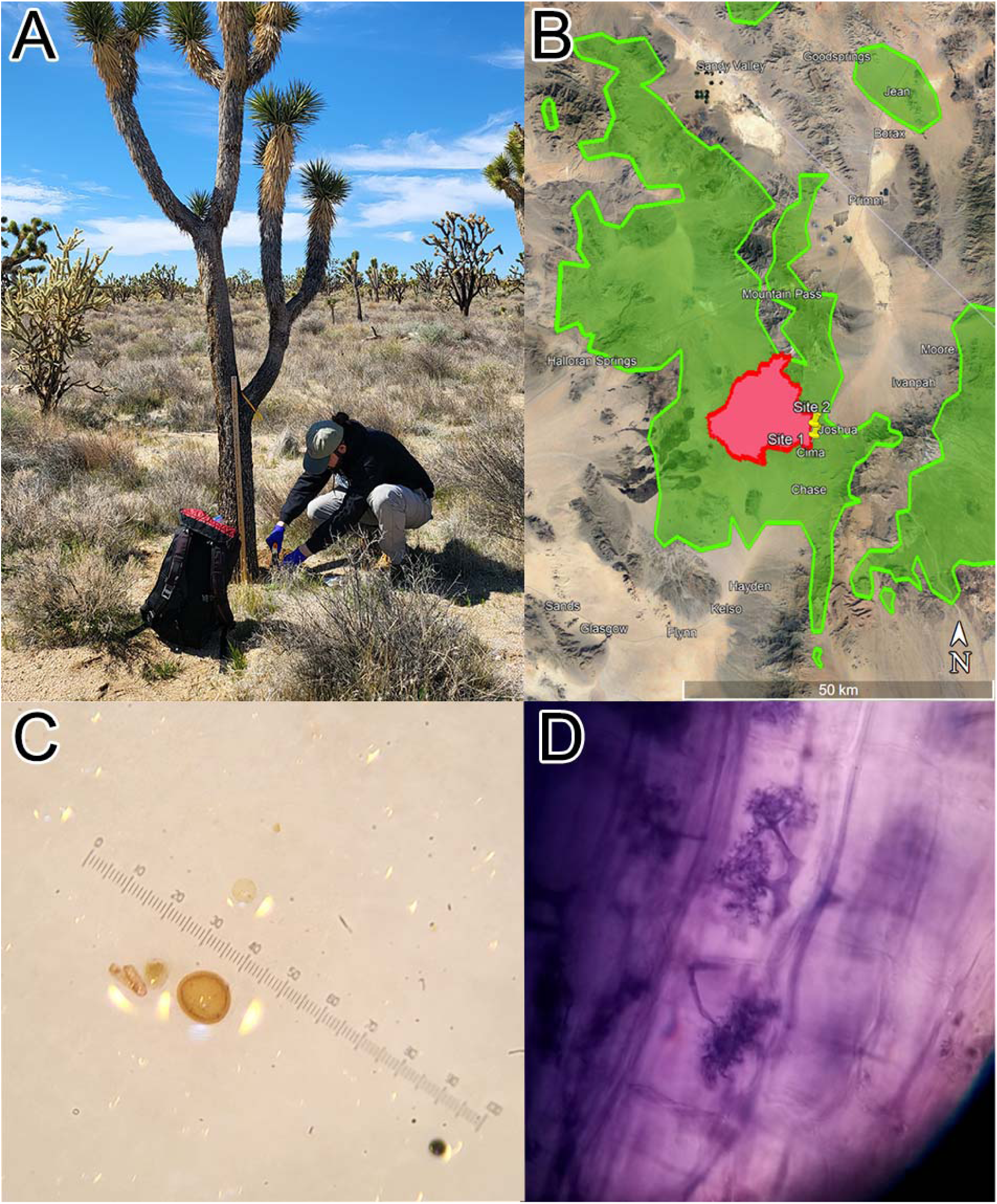
(A) Spring sampling of Y. jaegeriana soil and roots. (B) Y. jaegeriana habitat near the USA California-Nevada border (green). Location of the Dome Fire within California (red). Pins indicate general sampling sites for 20 trees. (C) AMF Spores at 65x magnification. (D) AMF arbuscules stained with Trypan Blue within Y. jaegeriana root.

### AMF Spore Count and Colonization Analysis

We extracted and counted spore number and estimated morphotype diversity from all 20 trees from all seasons. We used 25g of dried soil for sucrose-Calgon spore extractions (Ianson and Allen 1986) collected with 15mL distilled water.

We dispensed 1mL aliquots of well-mixed spore extracts onto filter paper and counted all spores at 40X magnification four times per sample. We averaged those 4 spore counts, multiplied by 15 (mL) to extrapolate the total count, and divided by 25 (g) to estimate spore content per gram of soil (Fig. 1C). We assessed spores for their relative size and color and assigned them to 12 different morphotypes (McKenney and Lindsey 1987). We confirmed the presence of AMF colonization in the roots (Fig. 1D) through trypan blue staining and microscopy (McGonigle et al. 1990). We rinsed roots in distilled water and cleared overnight in 2.5% KOH. We then rinsed roots in water again and bleached with 15% peroxide with 0.05% KOH for 5 minutes at room temperature to reduce red coloration from glycoside saponins in the roots.

We then acidified roots with 2% HCl for 2 minutes and moved roots to a 0.05% Trypan Blue solution at 90°C for 30 minutes. Finally, we destained the roots in a 1:1 solution of lactic acid and glycerol for 24 hours. We mounted roots on slides in PVLG to detect AMF structures but did not quantify colonization rates due to inconsistent clearance of red glycoside saponins from the roots.

### DNA Extractions

To extract DNA from soil, we used Qiagen DNeasy Powersoil Pro kits following the manufacturer’s protocol, except we replaced 100µL C1 solution with 100µL ATL buffer and incubated overnight at 4°C to improve yields. We followed an established protocol to extract DNA from roots (Glassman et al. 2015), where we lyophilized roots for 24 hours (Labconco FreeZone 4.5L, Kansas City, MO, USA), used sterilized steel beads to grind the roots using a 1 minute Fast-Prep 24™ (MP Biomedicals, Santa Ana, USA) cycle at 6.5m/sec, then added 1 ml of 1:1000 beta-mercaptoethanol:cetyltrimethyl ammonium bromide 2X (CTAB) and incubated one hour at 65°C. We then used 600µL of chloroform to remove lipids and mixed 700µL of the aqueous solution with 350µL 95% EtOH then proceeded with the Qiagen DNeasy Blood and Tissue Kit starting with buffer AW1.

### PCR and Sequencing

We used a two-step PCR to amplify the AMF-specific WANDA-AML2 primer pair (Lee et al. 2008; Dumbrell et al. 2011) adapting from established methods (Weber et al. 2019). We mixed 3 µL DNA from roots or soil with 0.5µL of 10 µM of each primer, 8.5 μL of Ultra-Pure Sterile Molecular Biology Grade water (Genesee Scientific), and 12.5 μl Accustart ToughMix (2× concentration; Quantabio). We amplified at the following settings: 94°C for 2 minutes, followed by 30 cycles of denaturing at 94°C for 30 s, annealing at 60°C for 30 s, elongating at 68°C for 45 s, and a final elongation step at 68°C for 2 minutes. We ran a 1.2% agarose gel to check for bands and increased to 35 cycles for samples with no visible bands. We cleaned PCR products with AMPure XP magnetic beads (Beckman Coulter Inc.) following manufacturer instructions. We used a second PCR to add DIP barcodes (Kozich et al. 2013) and adaptors for Illumina sequencing with a 94°C heating step for 2 minutes and 10 cycles at 94°C for 30 s, 60°C for 30 s, and 72°C for 1 min, using 12.5 µL Accustart Toughmix, 2.5µL of 1 µM DIP PCR2 primers, 6.5µL of ultra-pure water, and 1µL of PCR product. Following established protocols from our lab (Pulido-Chavez et al. 2023; Glassman et al. 2023), we pooled samples based on band intensity in an agarose gel by aliquoting 1 μL, 2 μL, or 3 μL per sample, then cleaned the pool with AMPure prior to checking concentration and quality with Agilent Bioanalyzer 210. We sequenced with Illumina Miseq 2x250 bp Nano kit, which provided sufficient sequence coverage due to low richness of AMF communities (Öpik et al. 2010), at the UC Riverside Institute for Integrative Genome Biology in 2 libraries to accommodate all samples. We included multiple negative controls for DNA extraction and PCRs in each library.

### Bioinformatics

We analyzed sequencing data using QIIME2 version 2022.2 (Bolyen et al. 2019). We removed forward and reverse adaptors using the QIIME2 cutadapt trim-paired plugin, demultiplexed, and denoised using DADA2 with the denoise-single plugin based on the quality of forward reads to ensure median quality scores over 30. We assigned taxonomy exclusively using forward reads, since the forward reads contain more of the informative variable region (Dumbrell et al. 2011; Davison et al. 2012). We matched sequences to the MaarjAM database (Öpik et al. 2010) using virtual taxa (VT) numbers and removed singletons to minimize the impact of erroneous reads. Since a portion of reads did not match to any VTs, they were subjected to a lower threshold BLAST against the MaarjAM database which retained matches with the highest e-value lower than 1e-50, a percent identity over 90%, and less than 10 nucleotide mismatches (Chaudhary et al., 2020). Sequences not matching one VT with these constraints were excluded from analysis as non-AMF reads.

### Statistical analysis

We performed statistical analysis in R version 4.1 (R Core Team 2020). We generated a species accumulation curve and rarefaction curve to determine cutoffs for sample numbers and sequencing depth using functions specaccum and rarecurve in “vegan” (Oksanen et al. 2022). We tested the impact of different rarefaction levels on number of samples, total number of VTs, and mean and median per sample AMF richness using the using the rrarefy.perm function in “vegan” and taking the mean of 10,000 permutations. We found that normalizing to 754 reads per sample allowed us to retain the most samples (143/150) and yield the same total number of VTs as non-rarefied data (47 VTs) but also account for uneven sequencing depth to accurately compare sample richness across treatments (Table S2). We calculated diversity indices using the estimateR and diversity functions in “vegan”.

To identify significant drivers of species richness across our trees, we used generalized linear mixed models from the “lme4” package (Bates et al. 2024). We used Poisson distributions after testing several distributions in the “MASS” package (Ripley et al. 2024) using the “car” (Fox et al. 2023) function qqp. We used reverse model selection to identify the lowest Akaike’s Information Criterion (AIC) value amongst differing generalized linear models incorporating season, type, and burn status with library and tree number held as random effects. We used chi-squared tests between reduced and full models when showing significance of season to test the impact of seasonality overall. To analyze seasonality of AMF VT richness and spore abundance, we used separate models for roots and soils due to differing levels of variance in the two groups. We confirmed normal distributions and low variance with the Shapiro-Wilk test and Levene’s test, respectively (Shapiro and Wilk 1965; Schultz 1985). Then we compared seasons with repeated measures ANOVA. We used the R package “stargazer” (Hlavac 2022) to export tables of statistics.

We analyzed differences in beta-diversity using Bray-Curtis dissimilarity matrices calculated with the “vegan” function avgdist rarefied to 754 sequences per sample, then applied a PERMANOVA analysis using the “vegan” function adonis2 and betadisper for differences between centroids and variation in mean distance of samples to centroids, respectively. We visualized beta diversity using non-metric multi-dimensional scaling (NMDS) plots with the “vegan” function metaMDS.

We generated masked sequence alignments of representative sequences of our forward read VTs and the Mucoromycota outgroup Endogone botryocarpus (Genbank accession LC431079). We made a phylogenetic tree of identified VTs using the QIIME2 plugin iq-tree-ultrafast- bootstrap which compared models to identify TIM+F+I+G4 (Transitional Model, Frequency, Invariant, rate 4 Gamma distribution) as the best fit model for tree construction based on the Bayesian Information Criterion.

We used heat maps and Venn diagrams to test and visualize the impact of sampling type (roots versus soils) and season on AMF richness and frequency and to identify a core microbiome. We used frequency of VTs in root and soil to generate Venn diagrams and made graphics using the “VennDiagram” package (Chen 2022). We used base R to create heatmaps based on prevalence of taxon appearance and selected color palettes from the “viridis” package (Garnier et al. 2024). We used the rankabundance function in the “BiodiversityR” package (Kindt 2024) to calculate the abundance and prevalence of VTs in our filtered and rarefied dataset.

Sequences were deposited to NCBI SRA PRJNA1196672. All scripts can be found at github.com/arik-chou/YJAMF.

## Experimental Procedures

After merging two Illumina libraries consisting of 2,399,441 reads, and filtering for quality, we obtained 1,264,130 sequences corresponding to 1,269 amplicon sequence variants (ASVs) from 150 samples, as 10 samples did not amplify. ASVs that matched a MaarjAM database VT were combined into 47 distinct VTs. Species saturation curves showed that sampling roughly 10 trees yielded most of the AMF VT diversity present (Fig. S1A). Rarefaction curves showed that all samples saturated at roughly 15 species per root and under 20 species per soil regardless of sequencing depth up ranging from 754 to 40,000 reads per sample (Fig. S1B).

### Spore abundance variation across seasons

Spore abundance averaged at a mean ± standard error (SE) of 12.06 ± 0.71 spores/1g soil, with 12 distinct morphotypes detected (Table S3). Summer had significantly fewer spores per gram than the other seasons (Table 1, Fig. 2A; (F_3,57_ = 10.3, p < 0.001). The Summer sampling (June 2021) followed the months with the lowest levels of precipitation (Fig. 2C). Spore abundance was driven by precipitation at each sampling time as a sum of the prior 3 months’ rainfall (F_1,76_ = 4.17, p < 0.0001, Table S4). Two morphotypes dominated in abundance (M3 and M4) across all seasons, followed by 2 more morphotypes (M5 and M8), which all had lowest abundances in the Summer with M4 and M8 peaking in Spring, M3 in Winter, and M5 in Fall. The remaining eight spore morphotypes showed similar abundance values, most peaking in abundance in Fall (Fig. S2).

**Figure 2.**
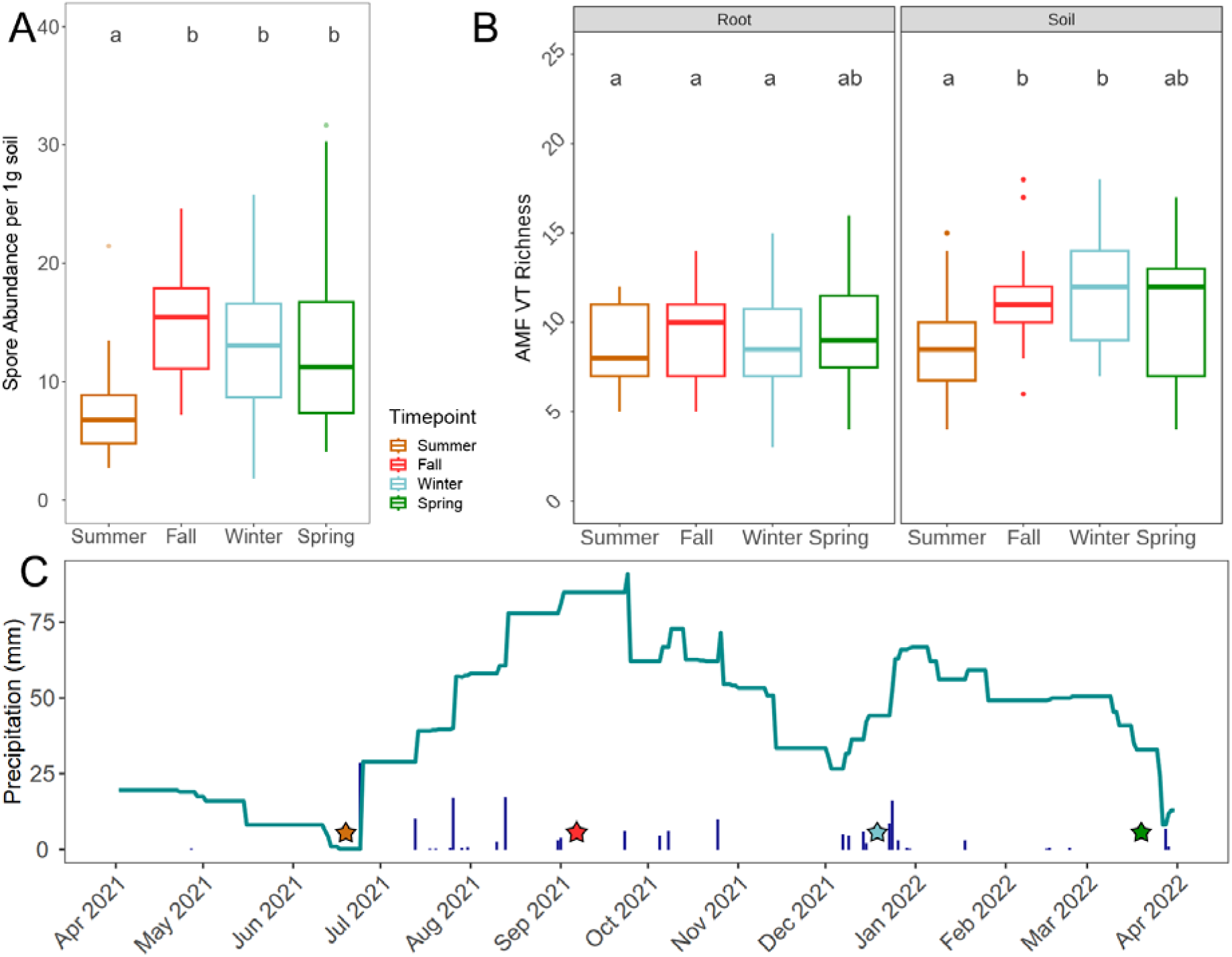
(A) Spore abundance in 1g of soil from 20 trees per season. (B) Richness of AMF virtual taxa in all trees per season in roots versus soils as detected by 18S sequencing. The median of each boxplot is indicated by the middle horizontal line, the bottom of the box indicates the first quartile, the top indicates the third quartile, and whiskers above and below indicate minimum and maximum values not considered outliers. Points beyond whiskers are outliers. Letters indicate statistically significant differences between least-squares means in our generalized mixed effects model. (C) Precipitation measured at the Mojave Mid Hills Station from April 2021 to March 2022. Bars indicate daily precipitation listed in mm, and the line indicates the sum of the prior 3 months of precipitation. Stars indicate our four seasonal sampling timepoints.

**Table 1.**
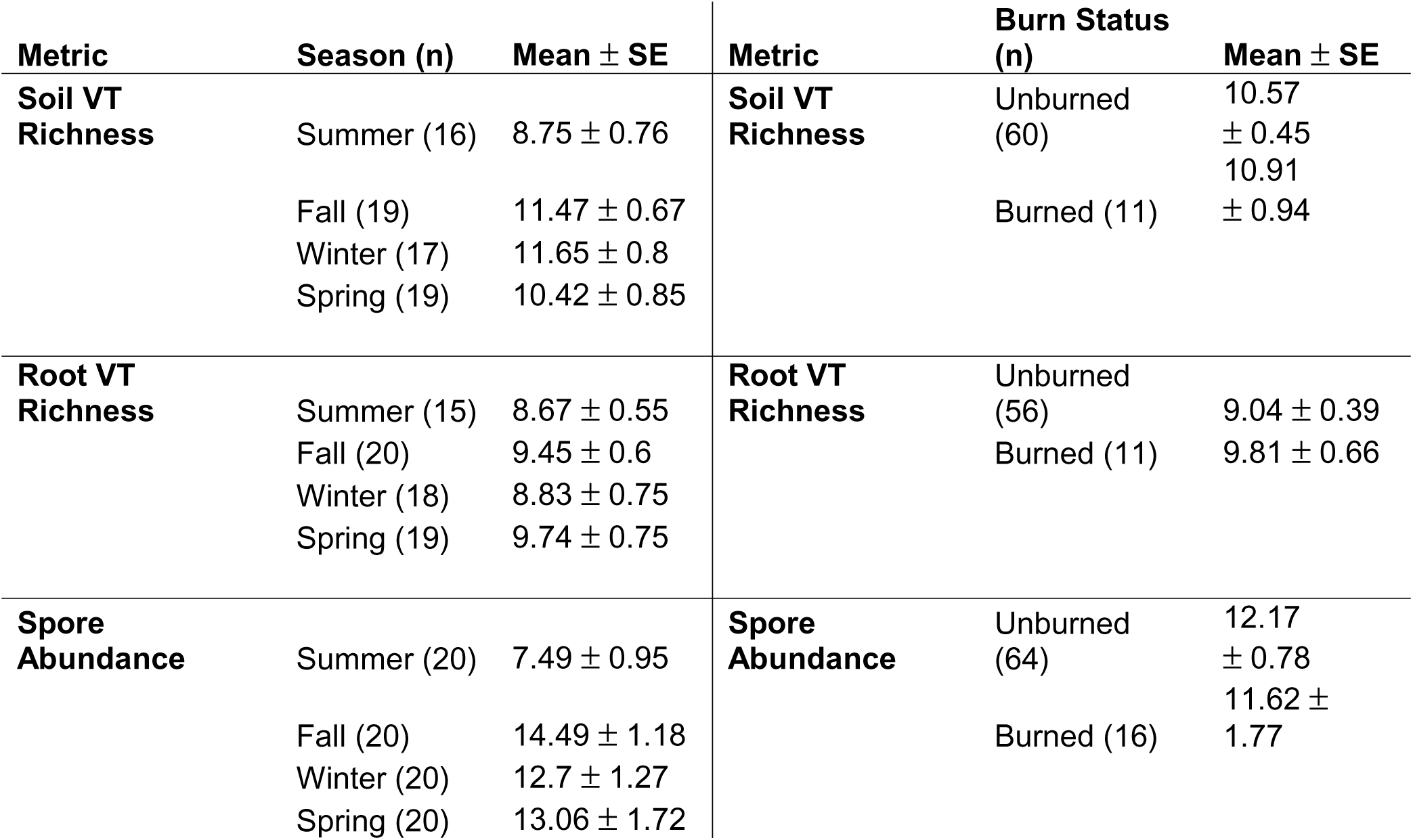
Per sample mean and standard error or AMF virtual taxa (VT) richness or spore abundance in different sample types, seasons, or burn statuses with the number of samples in parentheses.

| **Metric** | **Season (n)** | **Mean ± SE** | **Metric** | **Burn Status (n)** | **Mean ± SE** |
| --- | --- | --- | --- | --- | --- |
| **Soil VT Richness** | Summer (16) | 8.75 ± 0.76 | **Soil VT Richness** | Unburned (60) | 10.57 ± 0.45 |
| | Fall (19) | 11.47 ± 0.67 | | Burned (11) | 10.91 ± 0.94 |
| | Winter (17) | 11.65 ± 0.8 | | | |
| | Spring (19) | 10.42 ± 0.85 | | | |
| **Root VT Richness** | Summer (15) | 8.67 ± 0.55 | **Root VT Richness** | Unburned (56) | 9.04 ± 0.39 |
| | Fall (20) | 9.45 ± 0.6 | | Burned (11) | 9.81 ± 0.66 |
| | Winter (18) | 8.83 ± 0.75 | | | |
| | Spring (19) | 9.74 ± 0.75 | | | |
| **Spore Abundance** | Summer (20) | 7.49 ± 0.95 | **Spore Abundance** | Unburned (64) | 12.17 ± 0.78 |
| | Fall (20) | 14.49 ± 1.18 | | Burned (16) | 11.62 ± 1.77 |
| | Winter (20) | 12.7 ± 1.27 | | | |
| | Spring (20) | 13.06 ± 1.72 | | | |

### Impact of being in a burn patch on AMF richness and composition

Whether the tree was within the burn scar versus outside the burn scar did not significantly impact AMF VT richness (GLMM, z = 0.39, p = 0.70; Fig. 3A), so the burn effect was not included in our final model. There was also no significant difference in the mean spore abundance for trees within the burn scar versus outside the burn scar (Fig. 3B), but there was a small impact of burn on AMF VT community composition (PERMANOVA, F = 2.53, R^2^ = 0.018, p = 0.01).

**Figure 3.**
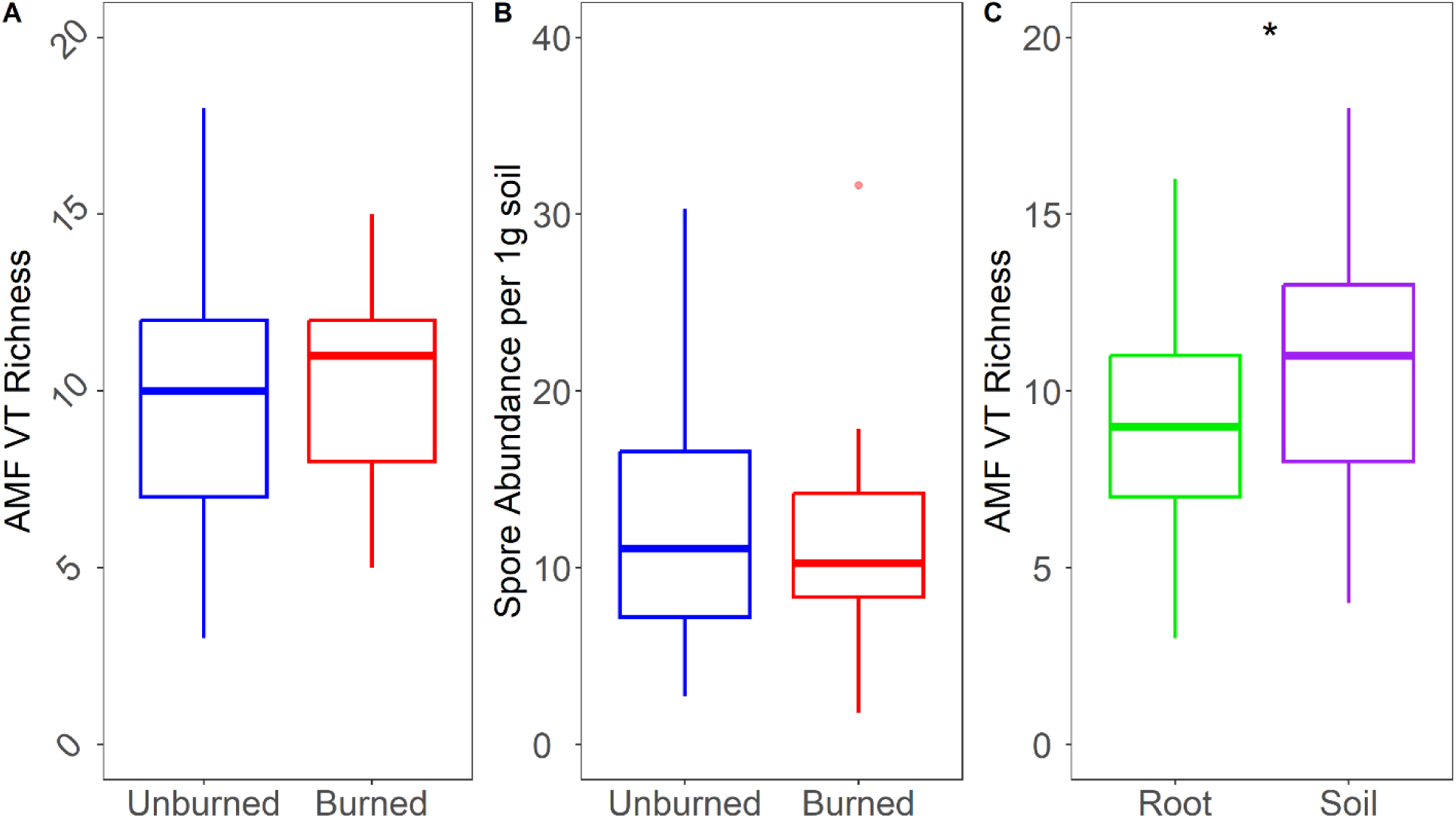
Comparison of burned and unburned Eastern Joshua trees and their AMF A) virtual taxa (VT) richness and B) spore abundance per 1g of soil. Blue represents unburned (n=16) and red represents burned (n=4) trees. Box plots show medians with the middle horizontal line, the bottom of the box indicating the first quartile, the top indicating the third quartile, and whiskers above and below indicating minimum and maximum values not considered outliers. Points beyond whiskers are outliers. No significant difference was found between either group. C) AMF Virtual taxa richness in root (n=71) and soil (n=72) samples combined across seasons. Stars indicate significantly greater richness in soil (p < 0.01).

### AMF richness and composition in roots versus soil

Averaged across seasons, soils yielded more AMF VTs per tree (10.63 ± 0.40 (n = 71)) than roots (9.18 ± 0.33 (n = 72)), (Fig. 3C, GLMM, z = 2.86, p = 0.004). Although soils yielded more AMF VTs than roots, 80% of AMF VTs were shared across soils and roots, with 38 of the 47 VT found in both sample types, and with 6 AMF VT exclusive to soil and only 3 to roots (Fig. 4A). Glomus VTX00294 was the most frequent AMF VT in both roots and soils (Fig. 5), detected in 87% of all samples. AMF VT communities showed small but significant differences between roots and soils in both the location of centroids (PERMANOVA, F = 15.0, R^2^ = 0.096, p = 0.001) and mean distance to centroids per sample (betadisper, F_1,1_ = 21.1, p < 0.001) when all seasons were considered together. Considering seasons separately, differences between AMF community composition of roots versus soils ranged from R^2^ of 0.07 in Summer and 0.08 in Spring to 0.13 in Fall and 0.19 in Winter (Fig. 6).

**Figure 4.**
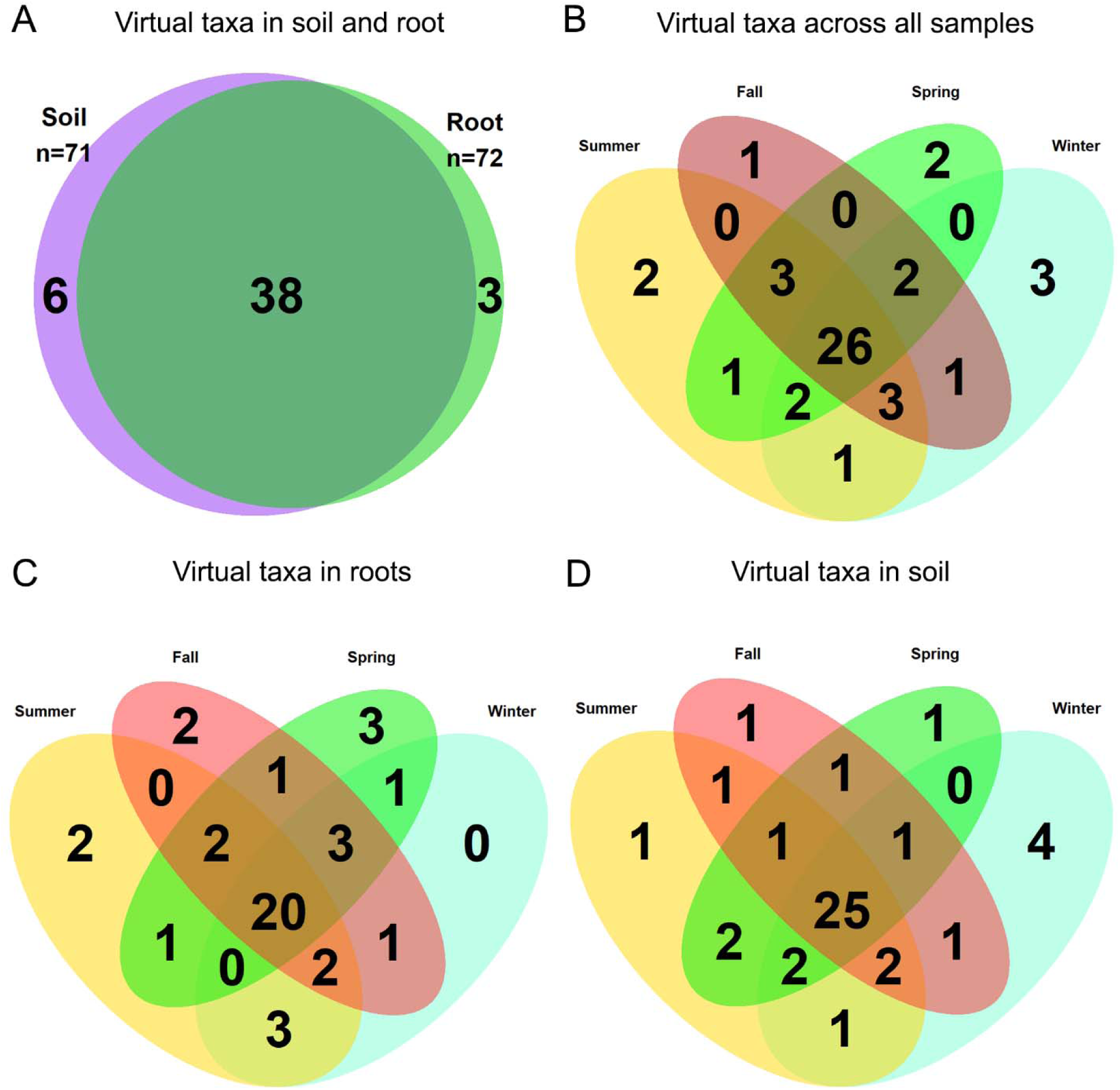
(A) Venn Diagram comparing virtual taxa in root and soil samples. (B) Venn Diagram indicating overlap of 47 detected virtual taxa in all root and soil samples. (C) Venn Diagram of 41 AMF VTs from roots and D) 44 soil VTs from each season.

**Figure 5.**
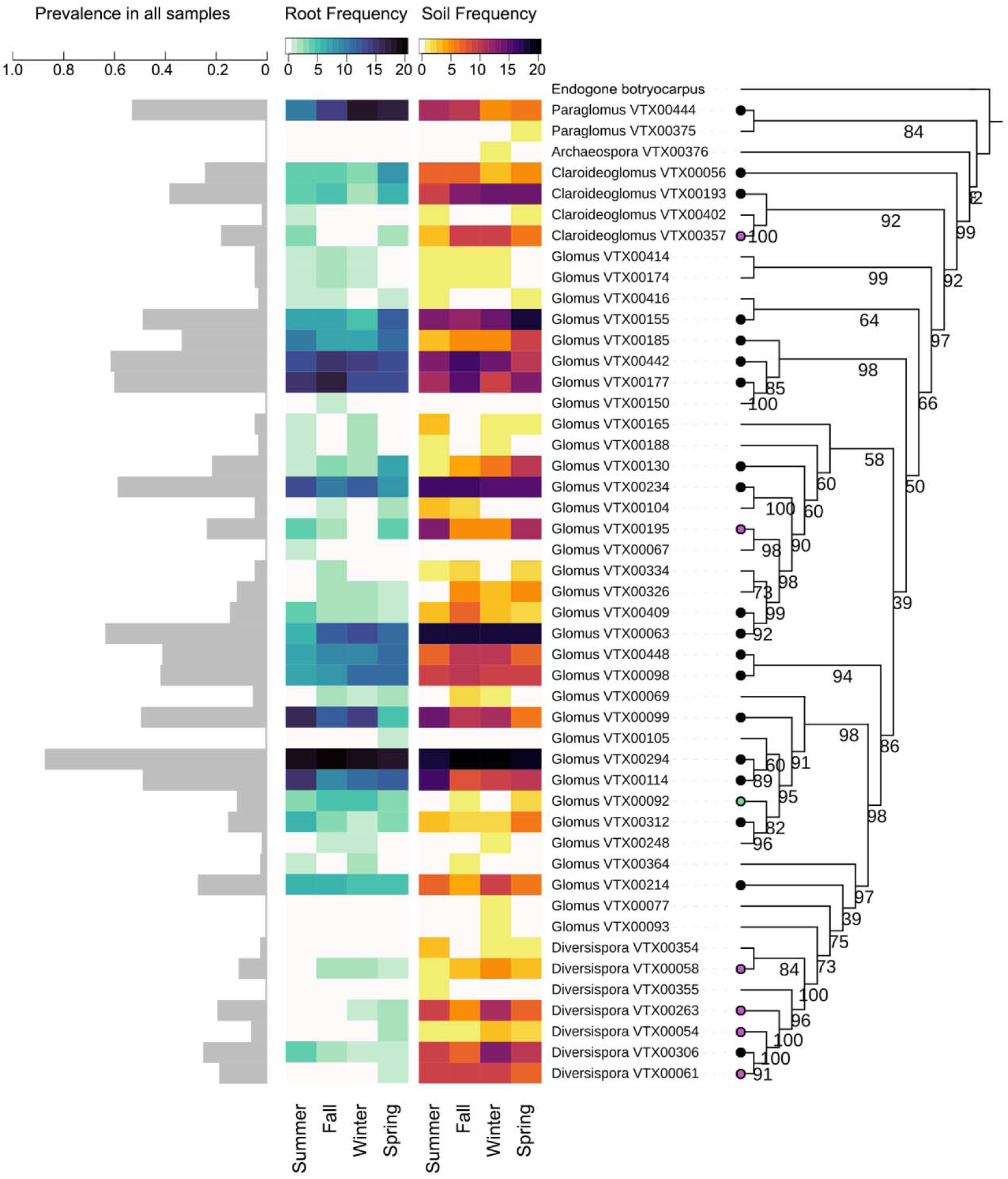
Prevalence of virtual taxa across 143 samples (gray). Heatmaps indicate AMF VT frequency across twenty trees in each season from roots (center-left) and soil (center-right). Root color scale in blue and soil color scale in red indicating frequency in up to 20 trees sampled per season. White cells are absences, and white rows indicate an absence of that VT from the sample type. A phylogenetic tree of 47 detected virtual taxa (with Endogone botryocarpus as an outgroup) was constructed based on forward reads. Numbers on the tree indicate bootstrap values. Tree-tip nodes indicate temporal core community members which are present in all 4 seasons in roots (green), in soils (purple), or in both (black).

**Figure 6.**
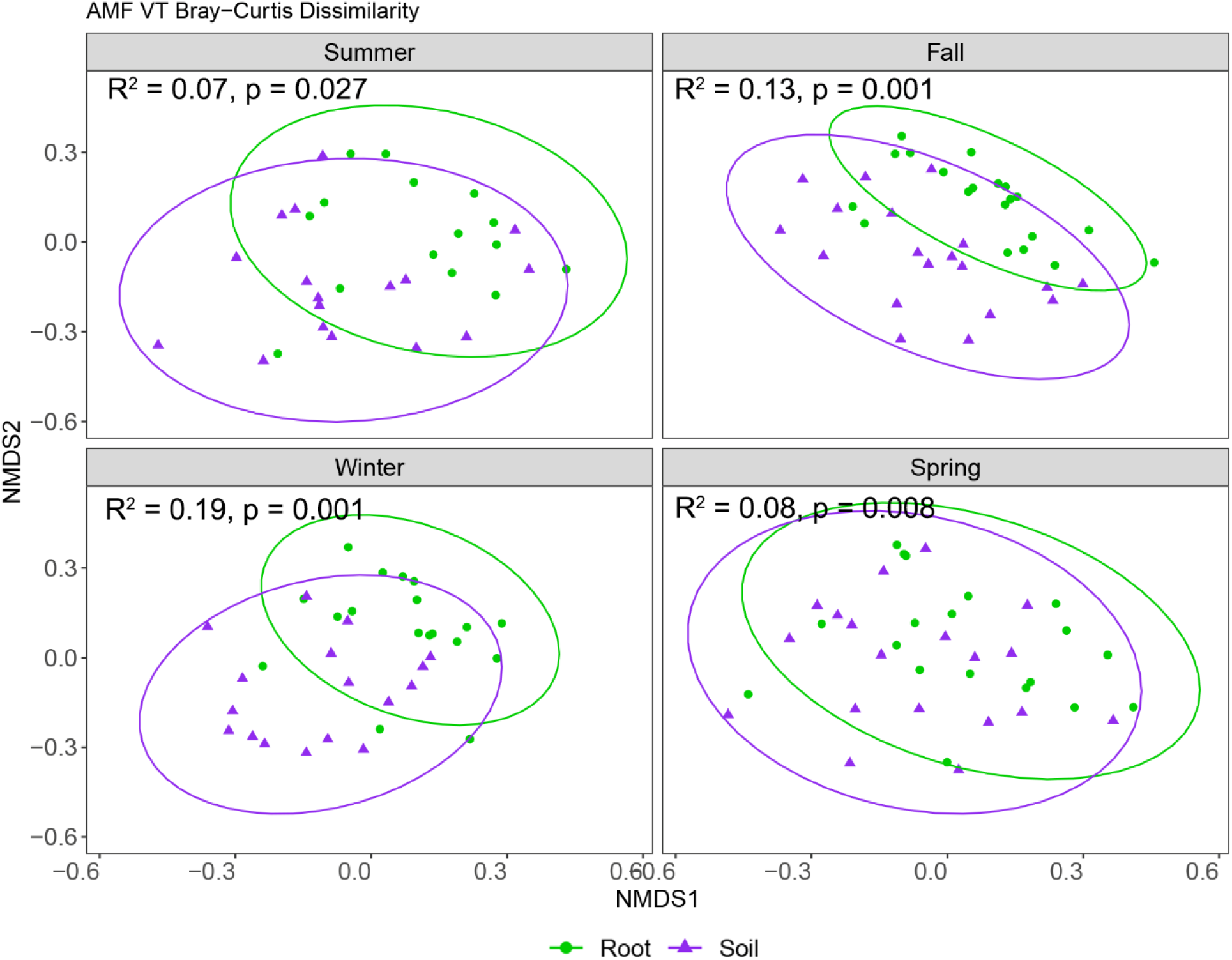
Bray-Curtis dissimilarity of AMF VTs across root (green circles) and soil (purple triangles) samples per season visualized as non-metric multidimensional scaling plots (k=3, stress = 0.18). Ellipses around each group represent 95% confidence intervals around the centroid of each sample type.

### AMF richness and composition across seasons

While there was no significant impact of season on per tree mean AMF richness in roots (F_3,46_ = 0.84, p = 0.48), soil samples did vary by season (F_3,45_ = 5.2, p = 0.004) with Summer having the lowest AMF VT richness (Table 1; Fig. 2B). Per tree mean AMF richness in soil was correlated with the sum of the prior 3 months’ rainfall in soil (GLMM, z = 3.03, p = 0.003) but not in roots (GLMM, z = 0.44, p = 0.66). When combining total per tree AMF richness from soil and roots, richness did not significantly vary by season (Fig. S3; F_3,44_ = 0.7, p = 0.6).

The majority (55%) of AMF VTs were found in all four seasons (Fig. 4B) and this pattern held for both roots (Fig. 4C) and soils (Fig. 4D). Seasons also had similar amounts of VTs detected overall when combining soil and roots, with 36 present in Fall and Spring, and 38 in Summer and Winter. However, each season had at least 1 VT that was not present in the other seasons (Fig. 4B), so season had a minor but significant impact on AMF community composition (Fig. S4, PERMANOVA, F = 2.00, R^2^ = 0.04, p = 0.004). In soil, community composition differed between Summer and Fall (F = 3.8, R^2^ = 0.10, p = 0.003), and Summer and Winter (F = 2.2, R^2^ = 0.06, p = 0.027). Meanwhile, roots differed only between Fall and Spring (F = 3.1, R^2^ = 0.07, p = 0.025), with no seasonal differences in beta-dispersion within root or soil. Heatmaps from root and soil samples (Fig. 5) highlight seasonal dominance for VTs highly prevalent across all samples, as the ten most prevalent VTs are also present in all seasons, representing 58% of the proportion of VT detection frequency. Some VTs were highly prevalent overall such as Glomus VTX00294 and Glomus VTX00063, and others had clear seasonal preferences for Summer (Glomus VTX00099 and Glomus VTX00114) or Fall (Glomus VTX00177 and Glomus VTX00442).

Sample-type specific trends included higher prevalence of Paraglomus VTX00444 in Summer soils, Diversispora VTX00306 in Summer roots, Glomus VTX00409 and Glomus VTX00098 in Fall soils, Glomus VTX00214 and Diversispora VTX00306 in Winter soils, and Claroideoglomus VTX00056 in Spring roots.

### Core community across roots/soil and seasons

A core community of 26 out of 47 VT primarily from the Glomeraceae was present across all seasons from roots and soils (Fig. 4B), with 20 out of 41 detected in roots (Fig. 4C) and 25 out of 44 in soil (Fig. 4D). A total of 19 VTs were present across all seasons in both roots and soils. Season-exclusive taxa included Glomus VTX00150 in Fall, Glomus VTX00067 and Diversispora VTX00355 in Summer, Glomus VTX00105 and Paraglomus VTX00375 in Spring, and Archaeospora VTX00375, Glomus VTX00093, and Glomus VTX00077 in Winter samples. While many of these exclusive taxa often only appeared in a few samples in one season with low overall prevalence (Fig. 5), other non-core community members appeared more frequently in other seasons but were completely absent in one season, such as Glomus VTX00069, Glomus VTX00165, Glomus VTX00174, Glomus VTX00414, Glomus VTX00416, and Glomus VTX00326. Of the 47 VT sequenced from roots and soil from Y. jaegeriana, only 9 VT overlapped with the 37 VTs associating with the closely related Y. brevifolia (Harrower and Gilbert 2021), those being Glomus VTX00114, Claroideoglomus VTX00193, Claroideoglomus VTX00056, Diversispora VTX00061, Diversispora VTX00306, Glomus VTX00092, Glomus VTX00069, Glomus VTX00093, and Glomus VTX00105, in descending order of frequency across all samples in this study. Of these, only Glomus VTX00114 showed high prevalence, as the eighth most prevalent taxa in our study.

## Discussion

Here, we present the first study of Eastern Joshua tree Yucca jaegeriana AMF and perform a comprehensive analysis of how AMF spore abundance, richness, and composition varies across seasons in roots versus soil among the same 20 trees sampled via both traditional microscopy and modern next generation sequencing methods. We confirmed AMF colonization in roots via staining and using Illumina MiSeq of WANDA-AML2 primers and identified 47 VTs overall from 5 families within the Glomeromycotina with a core AMF microbiome of 26 VTs across all seasons. AMF communities of Y. jaegeriana were diverse, with each tree harboring an average of ∼10 VTs per tree per sample type. Contrary to our predictions, AMF richness was unaffected by being within a burn scar, and roots did not yield greater AMF species richness than soil. In fact, soils yielded more AMF taxa than roots overall, although the vast majority of AMF VTs were shared by soil and roots. Further, AMF richness did not vary in roots across seasons but was lowest in Summer for soils and spore abundance. Notably, the AMF community was distinct from that of the sister species Y. brevifolia, with only 9 shared VTs between hosts, suggesting more host diversity than expected based on previous research indicating low host specificity among AMF (Davison et al. 2015).

We detected the lowest AMF spore abundance and richness in Summer soils following the driest months, but AMF spore abundance was constant from Fall to Spring. Similar trends regarding abundance and richness were reported in arid and semi-arid steppes (Bencherif et al. 2016), with richness and spore abundance at their lowest in the Summer. Precipitation leading up to the Summer sampling was minimal (Fig. 2C), likely slowing plant and AMF growth.

Sustained precipitation may have supported the increase in spore abundance we detected in Fall, similar to a study that detected a correlation between precipitation and spore density in semiarid shrublands (Chaudhary et al. 2014).

While we identified 4X as many AMF VT from next generation sequencing than from spore morphotyping, both followed similar trends. Overall richness of VTs and spore abundance were both lowest in summer and both showed high dominance by 1-2 taxa. Similar to our trends in VT prevalence, each of the four seasons represented a peak in abundance for at least one morphotype. This trend is consistent with prior studies in which spore abundance rose after precipitation when water is limiting, with some morphotypes still peaking in abundance outside of the most spore-dense season (Gemma et al. 1989; Bever et al. 2001; Da Costa et al. 2021; Kemmelmeier et al. 2022). While patterns of seasonality in spore morphotypes may be skewed by morphological plasticity of spores and other difficulties in species identification, a recent study of spore dispersal showed congruence between spore morphotyping and next-generation sequencing trends of aerial spores captured at a 20m height near agricultural fields (Chaudhary et al. 2020). While two morphotypes drove a peak in their post-harvest Summer spore counts, an overall diverse variety of 20 morphotypes and 17 VTs continued to follow their own seasonal trends with little change to VT richness, indicating that seasonal variation in spore dispersal is likely.

Eastern Joshua trees sampled within the burn scar did not show a decrease in AMF spore abundance or taxa richness, indicating that burning away the surrounding shrubs and grasses did not drastically impact the AMF associated with surviving trees. While we did not sample trees killed by the fire in this study, it is possible that spores would still be detected amongst the surrounding soil and dead roots, since AMF survive in storage after long periods of time (Allen 1992), and some burn conditions can even increase spore abundance (Aguilar-Fernández et al. 2009; Moura et al. 2022). Further, AMF beyond the top 10cm are thought to be buffered from low-intensity fires (Allen et al. 2011), thus seeding recolonization of shallow roots. While EMF appear to die off by 1 month post-fire (Pulido-Chavez et al. 2023), the longevity of AMF spores in the wild is not known. While it is possible that post-fire nutrient pulses reduce the need for AMF associations (Kennedy et al. 2002), other studies indicate that fast moving low intensity fires do not typically lead to long lasting impacts on MF richness (Chimal-Sánchez et al. 2015; Stürmer et al. 2022; Espinosa et al. 2023).

Most AMF taxa were shared by soil and roots, with AMF taxa exclusive to one sample type being rare. Indeed, 80% of AMF taxa were shared by soils and roots, and none of the sample-type exclusive taxa were found in any substantial frequency, often appearing in one sample. Six AMF VTs (Paraglomus VTX00375, Glomus VTX00093, Glomus VTX00077, Diversispora VTX00354, Diversispora VTX00355, and Archaeospora VTX00376) were detected in soil exclusively. These soil-exclusive taxa may have different colonization structures and strategies for soil or root, like Glomus intraradices and Glomus etunicatum (Hart and Reader, 2005). Further, Diversisporales AMF are well-known for their “edaphophilic” lifestyle of abundant extraradical hyphae and sparse hyphae in roots (Powell et al. 2009; Weber et al. 2019). This soil preference was supported by increased prevalence of Diversispora VTX00263 and Diversispora VTX00306 in Winter soils, but no corresponding increase of the same taxa in roots (Fig. 5). While some studies have shown substantial compositional differences in root and soil AM taxa (Hempel et al. 2007), we report significant overlap of root and soil communities with some variation in low abundance taxa.

Differing trends in seasonality across sample types led to some AMF compositional differences across seasons both between roots and soils and within roots and soils across seasons. For example, the differences between AMF composition in roots versus soils was higher in Fall and Winter (13-19%) compared to Summer and Spring (7-8%). Moreover, within soil AMF communities, they only differed between Summer-Fall and Summer-Winter, and within root AMF communities, they only differed between Fall-Spring but were consistent across other seasons. A study highlighting seasonal preference of AMF in nearby Palm desert surveyed multiple plant species’ roots to assess microbial responses to the Winter and Summer rains (Taniguchi et al. 2023). AMF taxa were a larger percentage of the fungal ITS1 sequence relative abundance after Summer rain but not Winter rains. Root AMF in wet-season months appeared to be structured based on associations with host plants rather than stochastic processes, consistent with community assembly patterns seen in sorghum fields after the removal of drought stress (Gao et al. 2020). Those patterns were reflected here in this study, as AMF VT richness in Spring was not significantly different from the lowest season, Summer, despite prior Winter rain (Fig. 2B). It is possible that ending the drought stress with Summer monsoons leads to more root growth and more opportunities for diverse AMF community structure in Fall. Another shallow-rooted succulent, Yucca angustissima, showed preferential Summer rain uptake while nearby perennials used both Summer and Winter rain sources (Ehleringer et al. 1991). Thus, root growth in Fall could support new root-specific mycorrhizal associations, while post-Winter rains do not, driving the compositional difference in Fall and Spring roots and resulting in differing amounts of variation across roots and soil.

AMF VT richness and community composition was otherwise largely consistent across seasons, with the majority of AMF VTs found across all seasons and with soil AMF richness reducing only in Summer. Our rarefaction curve shows that 10 Y. jaegeriana trees was sufficient for capturing most of the AMF community. Since we sampled double that, we were able to detect more rare taxa that either had some preference for roots versus soils or seasons, or were simply identified in one season versus another due to stochasticity. For example, Glomus VTX00104 was prevalent in Summer soils but not detected in Winter soils, but was detected in roots in the Spring. Other taxa share their peak in prevalence in both sample types at the same season, such as Glomus VTX00114 and Glomus VTX00099 in Summer, Glomus VTX00177 and Glomus VTX00442 in Fall, and Glomus VTX00155 in Spring.

We found 26 out of 47 VTs that were present in at least one root or soil sample from our 20 Y. jaegeriana trees at every season, forming a temporal core community. These include VTs from 4 of the 5 genera detected in this study including Glomus, Claroideoglomus, Paraglomus, and Diversispora ordered from most to least abundant. Whether these taxa are functionally critical or simply resilient is not known. It is possible that functional redundancies exist across the AMF community, but the high rate of genetic diversity across just one species of AMF suggests differing roles for each mycorrhizal association with implications for wider ecosystem functioning (Powell and Rillig 2018). Unlike other yuccas with deep taproots, Joshua tree roots are fibrous and shallow (Rundel and Gibson 1997), so they likely invest in a broad range of mycorrhizal partners to search for a variety of nutrients (Johnson 2010). This reliance on AMF may increase resource allocation towards many AMF species regardless of their benefit, potentially explaining the high diversity and low overlap of species between Y. jaegeriana and Y. brevifolia (Harrower and Gilbert 2021). Better understanding the dynamics of the core community will require investigations into differing contributions of AMF species to their plant host, as AMF communities often exhibit rare taxa appearing and disappearing with unclear consequences (Sánchez-Castro et al. 2012).

Surprisingly, despite their close genetic relationship and the general lack of host specificity among AMF (Davison et al. 2015), only 9 of 47 VTs overlapped between Y. jaegeriana and Y. brevifolia, even while targeting the same V3 portion of the fungal 18S rRNA and assigning taxonomy with the same MaarjAM database (Harrower and Gilbert, 2021). The most abundant VT that we found on Y. jaegeriana, Glomus VTX00294, has also been detected nearby in the Chihuahuan Desert in New Mexico (Porras-Alfaro et al. 2007), and as far as Australia (Davison et al. 2015) but not in Y. brevifolia. Expected genera that were detected in JTNP with Y. brevifolia such as Ambispora as well as Scutellospora were not present in the Cima Dome at their corresponding elevations from JTNP (Harrower and Gilbert 2021). This suggests that climatic or edaphic characteristics vary enough to support diverging AMF communities in closely related plants, potentially lending further support for host specificity of AMF (Kajihara et al. 2022; d’Entremont and Kivlin 2023). The distinct AMF VTs associating with each Yucca suggests that approaches using mycorrhizal inocula in restoration of Western and Eastern Joshua tree habitat should be independently developed.

## Conclusion

Although we focus narrowly on the AMF of a single desert tree, our comprehensive sampling across space, time, and roots versus soils mean that our results have implications both for restoration of endangered and iconic Joshua trees, and for general understanding of variation in time and space of host-associated microbiomes. We characterized for the first time the AMF the iconic Eastern Joshua tree and found that a single desert tree species associates with a diverse assemblage of 47 AMF taxa from 5 families within the Glomeromycotina. In addition, there is low overlap of AMF VTs shared between closely related sister genera of Yucca, indicating high levels of desert biodiversity are found belowground. Sampling soils is typically less effort than searching for roots especially in deep rooted desert plants, and our findings indicate that sampling from soils is preferred since most AMF taxa were shared but soils had higher richness and more unique VTs than roots. We also identified all seasons besides Summer as equally viable options for sampling, since we found high consistency in AMF spore abundance, richness, and composition across Fall, Winter, and Spring. Finally, our in-depth examination of both root and soil from the same 20 trees across seasons gives insight into generalized principles of host-associated microbiome turnover across space and time, indicating that a diverse core AMF microbiome persists across seasons and sampling type.

## Acknowledgments

We acknowledge Mojave National Preserve and the Mojave Desert Land Trust for allowing us to sample and Debra Hughson, Annasofia Andeski, and Andrew Kaiser for help with permitting. We thank Tasha La Doux and Jim Andre at the Sweeney Granite Mountains Desert Research Center for assistance with field sampling. We thank Edie and Mike Allen for their extensive help and training with methods for AMF spore extractions and root colonization and help with identification via spore morphology. We thank Sohrab Bodaghi, Gerardo Uribe, and Georgios Vidalakis for assistance and use of their lyophilizer. We thank Victoria Sloot and Justin Diab for their assistance with spore counting and identification. We thank Aishwarya Veerabahu, Marcos Vinicius Caiafa Sepulveda, Dylan Enright, Anna Nguyen, and Basubi Binti Zhilik for assistance with field work. We thank Joel Sachs and Quinn McFrederick for their input on the manuscript and the anonymous reviewers for their suggestions and revisions. We thank the Shipley Skinner Riverside County Endowment to AJ and SIG and the USDA Hatch fund #CA-R-MPP-5232-H to SIG for funding.

