## Supplemental Material for "Eastern Joshua tree arbuscular mycorrhizal fungal mycobiomes largely consistent across roots, soils, and seasons"

Supplementary Materials

Table S1 – GPS Location of Joshua trees sampled for root and soil.

| Tree Number | Latitude | Longitude | Height (cm) |
| --- | --- | --- | --- |
| 1 | 35.251275 | -115.497708 | 325 |
| 2 | 35.251261 | -115.49756 | 302 |
| 3 | 35.250905 | -115.497428 | 420 |
| 4 | 35.251377 | -115.497459 | 335 |
| 5 | 35.251398 | -115.497396 | 249 |
| 6 | 35.251412 | -115.497392 | 230 |
| 7 | 35.251411 | -115.497686 | 422 |
| 8 | 35.25169 | -115.497233 | 567 |
| 9 | 35.268193 | -115.498134 | 577 |
| 10 | 35.268291 | -115.498224 | 308 |
| 11 | 35.267998 | -115.497597 | 434 |
| 12 | 35.268044 | -115.497910 | 506 |
| 13 | 35.267713 | -115.497311 | 281 |
| 14 | 35.268048 | -115.497525 | 588 |
| 15 | 35.267264 | -115.497235 | 356 |
| 16 | 35.267647 | -115.497137 | 272 |
| 17 | 35.250607 | -115.502029 | 402 |
| 18 | 35.250872 | -115.501821 | 343 |
| 19 | 35.251253 | -115.501689 | 173 |
| 20 | 35.250872 | -115.501821 | 294 |

Table S2. Sequence rarefaction levels of our sample-VT table of 150 samples and 47 virtual taxa. Prior to rarefaction, each sample had a median 5,565 reads and median 10 VTs. Low rarefaction levels were tested to maintain sample numbers and VTs. Increasing rarefaction to over 1,000 sequence reads per sample removed samples below that threshold which were low in species abundance, increasing the mean species richness. Thus 754 reads were selected for maintaining the total VT count and preserving the most sample numbers.

|  | Mean Species | Median Species | Mean Seq | Median Seq | Total Taxa | N (samples) |
| --- | --- | --- | --- | --- | --- | --- |
| **Pre-rarefaction** | 10.1 | 10 | 7266 | 5565 | 47 | 150 |
| 418 Reads | 9.6 | 9 | 418 | 418 | 45 | 146 |
| 754 Reads | 9.9 | 10 | 754 | 754 | 47 | 145 |
| 889 Reads | 9.9 | 10 | 899 | 899 | 47 | 143 |
| 10% -  1488 Reads | 10.3 | 10 | 1488 | 1488 | 47 | 135 |

Table S3. Spore morphotypes based on single spore photos measured for identification purposes. Potential identifications done by expert opinion from Dr. Michael Allen.

| Type | Size | Color | Note | Potential Taxon |
| --- | --- | --- | --- | --- |
| M1 | 500 µm | Hyaline |  | *Gigaspora* |
| M2 | 250 µm | Orange-red |  | *Acaulospora* |
| M3 | 24 µm | Hyaline |  | *Glomus mosseae* |
| M4 | 24 µm | Yellow-orange |  | *Glomus* |
| M5 | 10 µm | Yellow to hyaline | Clustered | *Glomus aggregatum* |
| M6 | 85 µm | Yellow | Clustered | *Glomus* |
| M7 | 200 µm | Hyaline | 1-2 spots within spore | *Scutellospora* |
| M8 | 10 µm | Red to purple |  |  |
| M9 | 250 µm | Yellow | Budding |  |
| M10 | 20 µm | Red to purple | Clustered |  |
| M11 | 50 µm | Hyaline with red inside |  |  |
| M12 | 300 µm | Translucent brown |  |  |

Table S4. A comparison of generalized mixed effects models of spore abundance, comparing a model with Fall, Winter, and Spring to the Summer reference level (Model 1), a model replacing seasons with a sum of the prior 3 months’ precipitation (Model 2), and a model with a sum of 1 years’ precipitation (Model 3). Precipitation values were scaled around zero. Each row lists the estimated coefficient and standard error in parentheses beneath it. Significance is indicated by stars. Both models use root and Summer as the Sample type and Season reference value, respectively.

| *Dependent variable:* | | |  |
| --- | --- | --- | --- |
|  | Spores per 1g | | |
|  | (1) | (2) | (3) |
| TimepointFall | 0.749^***^ |  |  |
|  | (0.004) |  |  |
| TimepointWinter | 0.564^***^ |  |  |
|  | (0.004) |  |  |
| TimepointSpring | 0.567^***^ |  |  |
|  | (0.004) |  |  |
| Sum of Prior  3 Months’ Rain |  | 0.209^***^ |  |
|  |  | (0.050) |  |
| Sum of Prior  1 Years’ Rain |  |  | 0.277^***^ |
|  |  |  | (0.004) |
| Constant | 1.932^***^ | 2.420^***^ | 2.405^***^ |
|  | (0.004) | (0.091) | (0.004) |
| Observations | 80 | 80 | 80 |
| Log Likelihood | -237.235 | -244.315 | -238.204 |
| Akaike Inf. Crit. | 486.471 | 496.629 | 484.408 |
| Bayesian Inf. Crit. | 500.763 | 506.157 | 493.936 |
| *Note:* | ^*^p<0.1; ^**^p<0.05; ^***^p<0.01 | | |

Table S5. Models of seasonality on VT richness in soil and root samples. Seasons drove significant differences in soil but not root samples. Each row lists the estimated coefficient and standard error in parentheses beneath it. Significance is indicated by stars. Summer is used as the reference level for comparison.

| *Dependent variable: VT Richness* | |  |
| --- | --- | --- |
|  | (Soil) | (Root) |
| TimepointFall | 2.868^***^ | 0.897 |
|  | (0.832) | (0.823) |
| TimepointWinter | 3.010^***^ | 0.369 |
|  | (0.890) | (0.811) |
| TimepointSpring | 1.787^**^ | 1.186 |
|  | (0.870) | (0.807) |
| Constant | 8.680^***^ | 8.553^***^ |
|  | (0.789) | (0.724) |
| Observations | 71 | 72 |
| Log Likelihood | -173.272 | -169.837 |
| Akaike Inf. Crit. | 360.543 | 353.675 |
| Bayesian Inf. Crit. | 375.976 | 369.211 |
| *Note:* | ^*^p<0.1; ^**^p<0.05; ^***^p<0.01 | |

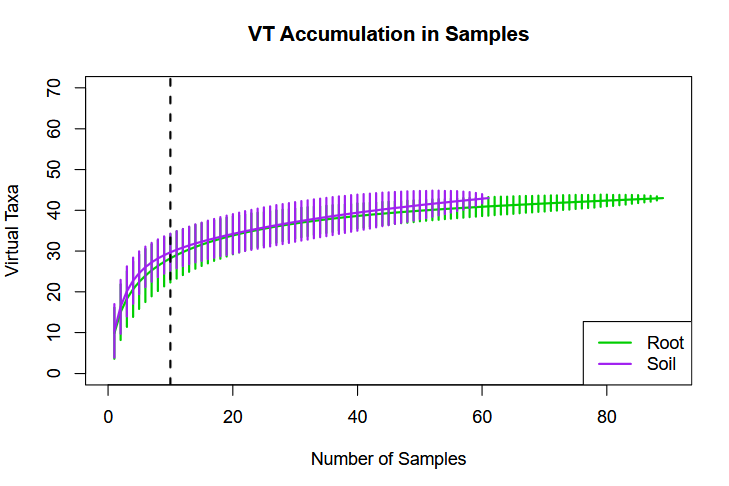

**A**

**B**

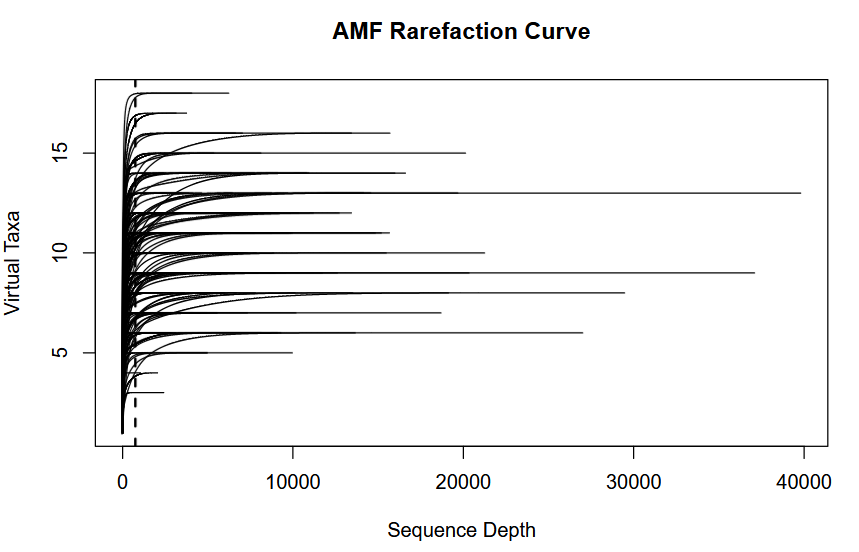

Figure S1. A) Species saturation curves based on 47 virtual taxa detected in all our samples across 1 year. Soil samples are depicted as purple and root samples are depicted in green. The curve indicates how many samples are sufficient for capturing most of the species present in the community. The dashed line at 10 samples shows approximately where taxa accumulation levels off. B) Sequence depth compared to the amount of species detected. Each line is one sample. Our rarefaction level of 754 sequences (dashed line) allows most samples to saturate and importantly, preserves the total number of VT detected without removing any.

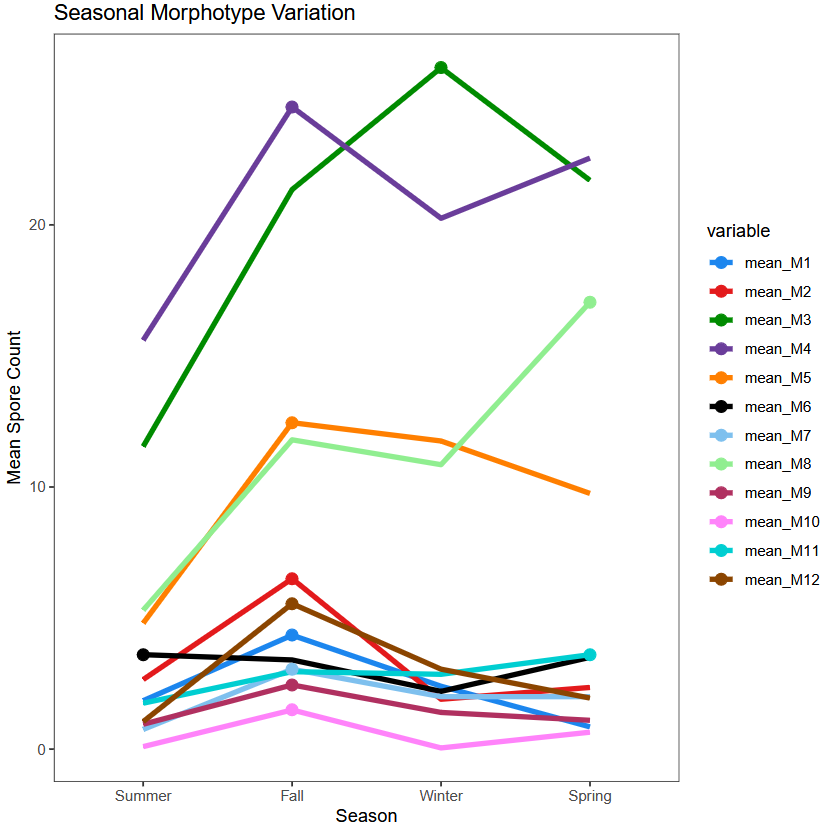

Figure S2. Mean spore morphotype count in twenty trees across each season. Circles indicate the season in which each morphotype was most abundant.

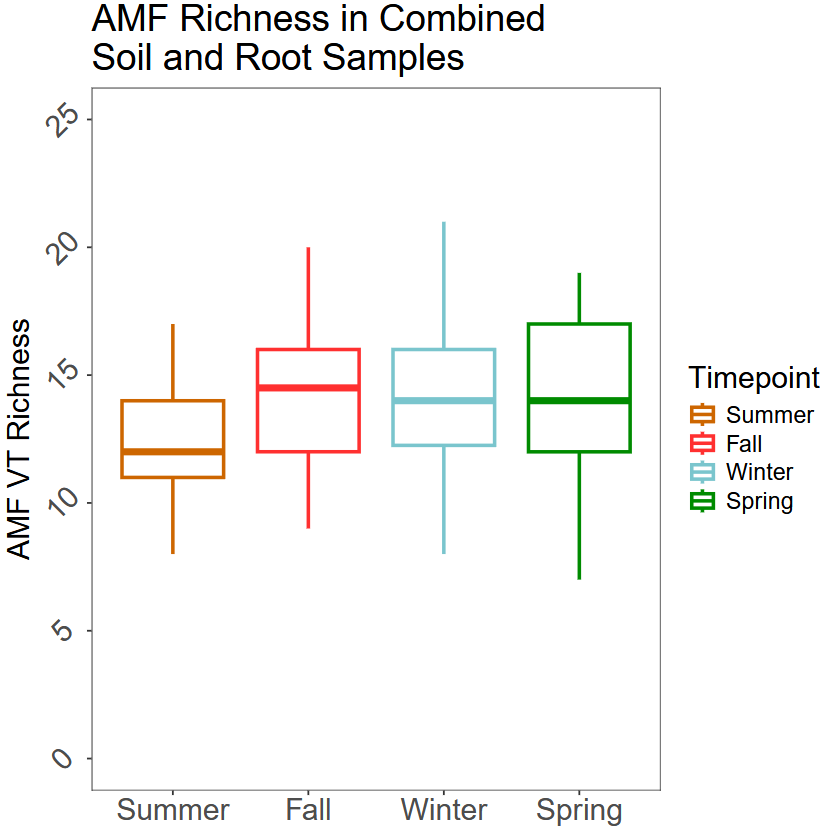

Figure S3. Richness comparisons of each season (from both root and soil combined per tree). Box plots show medians with the middle horizontal line, the bottom of the box indicating the first quartile, the top indicating the third quartile, and whiskers above and below indicating minimum and maximum values not considered outliers.

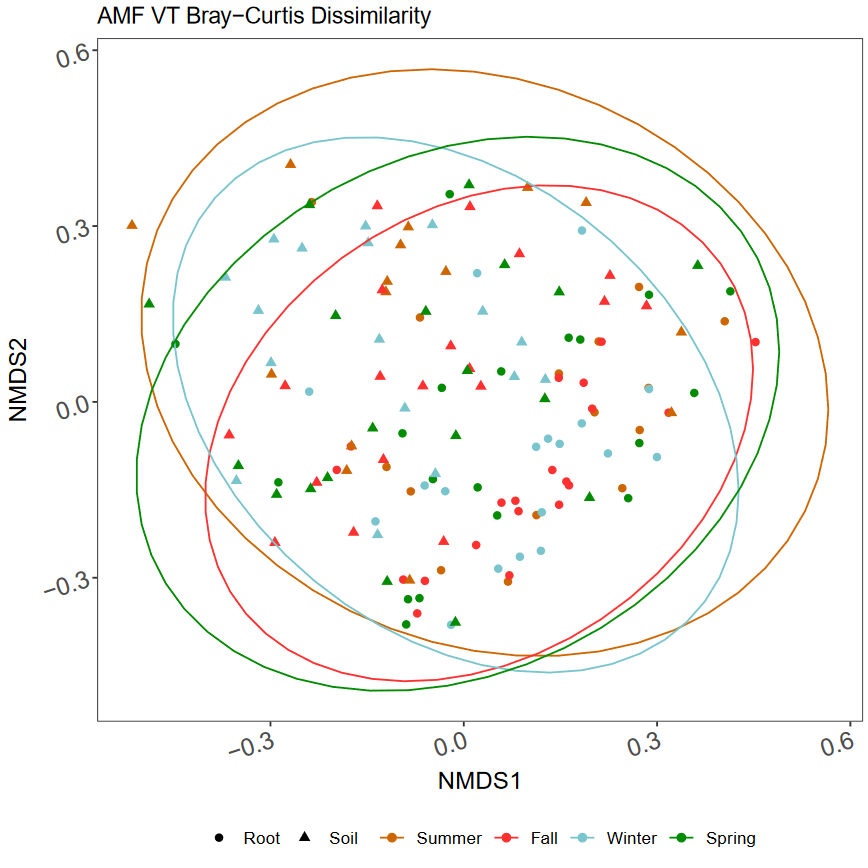

Figure S4. Bray-Curtis dissimilarity of samples based on VT sequences across seasons visualized as non-metric multidimensional scaling plots (k = 3, stress = 0.18). Triangles indicate soil samples and circles indicate root samples.
